# Ultra-High Resolution Solid-State NMR for High Molecular Weight Proteins on GHz-Class Spectrometers

**DOI:** 10.1101/2025.05.05.652283

**Authors:** Songlin Wang, Thirupathi Ravula, John A. Stringer, Peter L. Gor’kov, Owen A. Warmuth, Christopher G. Williams, Alex F. Thome, Leonard J. Mueller, Chad M. Rienstra

## Abstract

NMR spectroscopy is a powerful technique with broad impact across the physical and life sciences, and ultra-high field, GHz-class NMR spectrometers offer exceptional overall performance including superior resolution and sensitivity. While the resolution is fundamentally limited by molecular tumbling for solution NMR, solid-state NMR (SSNMR) is constrained only by instrumentation, making it well-suited for studying large and complex systems. To fully leverage UHF magnets for magic-angle-spinning SSNMR, it is essential to eliminate linebroadening arising from magnetic field drift and couplings among the nuclear spins. We address these challenges using an external ^2^H lock to compensate the field drift and Long-Observation-Window Band-Selective Homonuclear Decoupling (LOW-BASHD) to suppress ^13^C homonuclear couplings. We thereby achieve better than 0.2 ppm resolution in proteins up to 144 kDa, enabling unique site resolution for over 500 amide backbone pairs in 2D experiments. This exceeds the resolution available from solution NMR for large biological molecules, greatly expanding the potential of GHz-class NMR for research in life sciences.

**Teaser:** Ultra-high field NMR enables scientists to observe the finest details of large biomolecules, and this study overcomes key challenges to achieve a new benchmark for resolution in solid-state NMR of high molecular weight proteins.

## Introduction

With advancements in superconducting technologies, nuclear magnetic resonance (NMR) has entered the GHz (10^9^ Hz) era. Ultra-high field (UHF) NMR spectrometers now offer outstanding resolution and sensitivity, substantially facilitating research in materials and life sciences.(*1-4*) High molecular weight biological systems, such as enzymes, assemblies, transporters, receptors, and motor proteins, particularly benefit from increased field strength, as the improved resolution enhances spectral dispersion and enables the isolation of signals of interest from overlapping resonance peaks. However, solution NMR faces inherently limitations in studying large biological molecules because its resolution is constrained by molecular tumbling rates. As molecular size increases, slower tumbling leads to severe line broadening, diminishing the resolution gains from UHF. It restricts routine solution NMR studies to proteins below 40 kDa, except in a few special cases.(*5, 6*) In contrast, for magic-angle-spinning (MAS) solid-state NMR (SSNMR), resolution continues to improve with increasing magnetic field strength, regardless of molecular size. As a result, SSNMR is particularly well-suited for studying large and complex biological systems and benefits the most from advancements in magnet technology.

The development of high-field NMR has historically been constrained by magnet technology, with early instruments relying on resistive magnets that exhibited insufficient field homogeneity and temporal stability.(*7*) A breakthrough occurred in the 1960s with the introduction of NbTi superconducting magnets, which enabled field strengths up to approximately 270 MHz (6.3 T). At that time, concerns were raised regarding the feasibility of higher fields due to limitations in probe design and radiofrequency (RF) performance. However, Hoult and Richards provided a critical theoretical analysis demonstrating that increased magnetic fields would enhance sensitivity and resolution, overcoming initial skepticism.(*8*) Despite these advancements, NbTi-based magnets reached a practical limit of ∼400 MHz (9.4 T), leading to a decade-long stagnation in field progression.(*9*) The emergence of Nb□Sn superconducting wire and improved cryogenic technology in the late 1980s enabled the development of 500 and 600 MHz systems, followed by 750 MHz magnets by the mid-1990s. Further refinements in Nb□Sn processing, combined with the use of 2 K cryogenic pumping, allowed for 800 and 900 MHz systems by the early 2000s.(*10*) However, for nearly two decades, 900 MHz remained the practical upper limit of commercial NMR systems. A few specialized projects exceeded this threshold, such as the first GHz NMR (1.02 GHz) developed in Japan, and the 1.5 GHz Series Connected Hybrid (SCH) magnet at the National High Magnetic Field Laboratory (NHMFL), but these systems faced significant challenges, particularly regarding magnet stability and widespread accessibility.(*11, 12*)

The advent of high-temperature superconducting (HTS) materials, such as YBCO, has now overcome this limitation, enabling commercial GHz-class NMR spectrometers with field strengths of 1.1 and 1.2 GHz since 2020. Even higher fields, including 1.3 GHz, are currently in development.(*13, 14*) According to Bruker Corporation, more than 20 GHz-class NMR spectrometers have been installed globally over the past five years, and the number continues to grow rapidly. GHz-class NMR spectrometers provide outstanding resolution and sensitivity; however, these systems face challenges related to magnetic field instability, including rapid drift, non-linear fluctuations, and field gradients that depend upon the magnet bore temperature. These issues are particularly problematic in newly energized magnets and can persist for months or even years after installation, as a consequence of the unique properties of the HTS technology.(*15*) These instabilities degrade spectral quality and complicate the standardization of chemical shift referencing,(*16*) resulting in greater uncertainty in peak positions and compromising one of the major benefits of utilizing higher magnetic fields. While the internal deuterium (^2^H) lock technique has been widely employed in solution NMR to address field instability, its application in SSNMR has been limited, as internal locks cause degradation of the RF probe performance and many SSNMR samples lack ^2^H nuclei with sufficiently narrow signals to use for the lock control. Although examples of effective external ^2^H locks for MAS SSNMR have been demonstrated,(*17, 18*) SSNMR probes delivered along with the GHz-class magnets from Bruker lack this capability. One alternative approach utilizes simultaneous acquisition of reference spectra to reduce the variations when processing the data,(*19*) although this method requires modifications to standard data collection and processing workflows and may be limited to samples containing a strong, narrow solvent signal. Therefore, many types of samples, such as membrane proteins, fibrils, drug formulations, and complexes, are not compatible with this strategy.

In this study, we introduce SSNMR probes equipped with external ^2^H lock coils compatible with the unique geometrical limitations of UHF HTS magnets. We achieve sufficient resolution for high molecular weight microcrystalline protein samples to resolve the fine structure arising from homonuclear ^13^C-^13^C scalar couplings. We then apply Long-Observation-Window Band-Selective Homonuclear Decoupling (LOW-BASHD)(*20*) to enhance the resolution and sensitivity by a factor of two. In combination, these techniques yield ultra-high resolution ^13^C SSNMR spectra, with typical linewidths of 0.1-0.3 ppm for ^13^C and ^15^N in very large proteins, enabling the unique identification of the large majority of the 665 backbone amide pairs for a 144 kDa protein assembly in standard 2D spectra.

## Results

### Design of the SSNMR probe with external ^2^H lock

The design of a SSNMR probe with external ^2^H lock requires that both the sample and lock coils lie within the limited range of the homogeneous magnetic field (“sweet spot”) in HTS magnets. Whereas conventional magnets typically have a sweet spot of at least 60 mm, this specification is not achieved in the UHF magnets. To image the field gradient for the 25.8 Tesla (1.1 GHz ^1^H Larmor frequency) Bruker Ascend HTS magnet that was installed at the National Magnetic Resonance Facility at Madison (NMRFAM, University of Wisconsin-Madison) in late 2023, we varied the vertical position of a Phoenix 1.6 mm probe within the magnet, and acquired a series of ^2^H 1D spectra from D_2_O packed in a 1.6 mm rotor with spinning at 5.8 kHz. During this measurement, all shim parameters were set to zero to visualize coil images, which became narrower as the sample approached the magnetic field center (Fig. 1A). The position defined as 0 mm corresponds to the highest probe position allowed by the top of the shim stack. The magnetic field center was determined to be 20 mm below this point, with an effective field range of approximately 35 mm. Compared with conventional superconducting magnets, this geometry is approximately a factor of two smaller in the Z direction, requiring a significantly more compact design to accommodate the high-power sample coil as well as the lock coil within the same magnetic field.

**Fig. 1.**
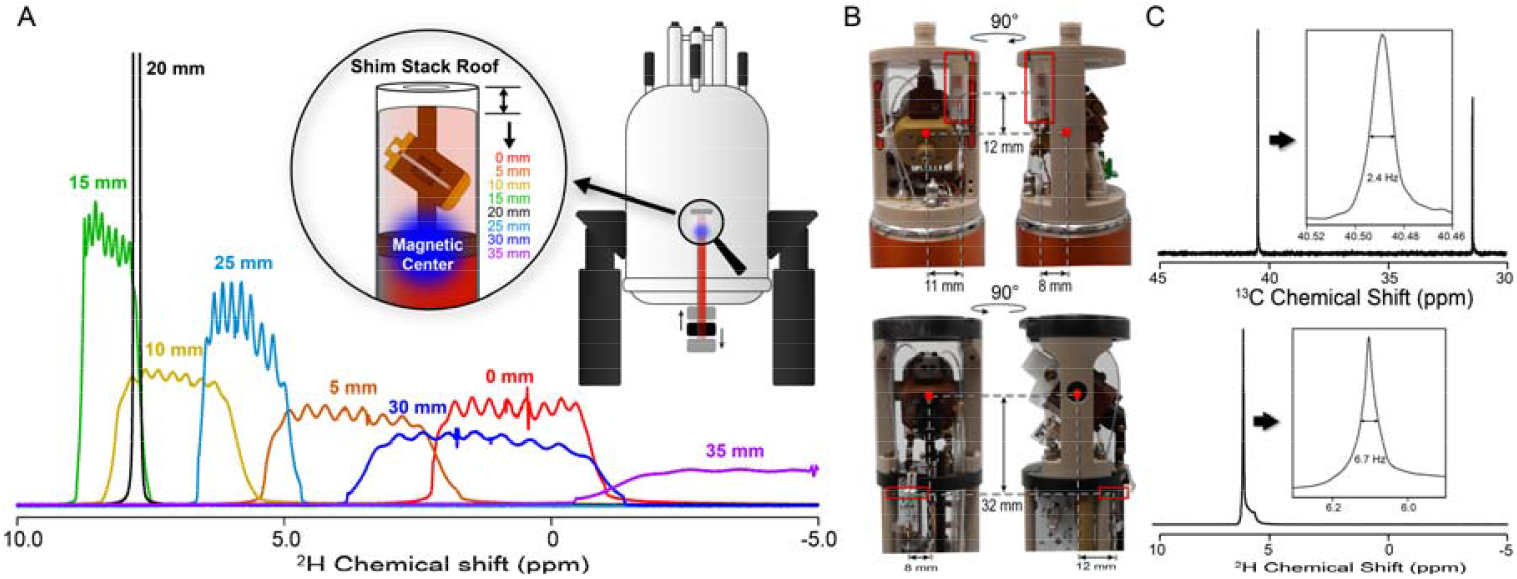
Mapping the effective field range and probe design with external ^2^H lock. (A) mapping the effective field range of the 1.1 GHz NMR spectrometer at NMRFAM. The positions are defined as distances to the highest probe position allowed by the top of the shim stack. (B) SSNMR probes with external ^2^H lock coil from PhoenixNMR (top) or Black Fox (bottom). Red dots indicate the centers of the sample coils, and red boxes highlight the ^2^H lock coils. The numbers illustrate the distances from the center of the sample coil to the center of the lock coil. (C) Optimal shimming results for adamantane in the sample coil (top) and D_2_O in the ^2^H lock coil (bottom) for the Black Fox 1.6 mm probe.

Fig. 1B illustrates two probes with suitable external ^2^H lock coils, incorporating epoxy sealed capillaries containing D_2_O as the lock sample. The vertical difference between the two coils is 12 mm for the Phoenix probe (Fig. 1B, top) and 32 mm for the Black Fox probe (Fig. 1B, bottom). To achieve the best shimming results for both coils, we detected ^13^C signal from adamantane in the sample coil(*21*) and ^2^H signal from D_2_O in the lock coil, using the Bruker NEO console capability for simultaneous acquisition with multiple receivers and a dual acquisition pulse program. We then shimmed the consensus magnetic field using a combination of automated and manual adjustments.(*22*) The Black Fox probe achieved linewidths of 2.6 Hz (10 ppb) for ^13^C and 6.7 Hz (40 ppb) for ^2^H (Fig.1C), with comparable performance observed for the Phoenix probe (Fig. S1). The probe heads utilize susceptibility matched materials and 3D modeling to symmetrize the components near the sample to achieve ppb-scale homogeneity,(*23*) despite the high magnetic field. We determined that a stable lock signal could be obtained with ^2^H linewidths as large as 20 Hz without noticeable splitting, with a best practice of 10 Hz or less.

### Post-installation field drift of GHz-class NMR and performance of the external ^2^H lock

Relative to conventional, metal alloy superconducting wire, the initial field drift rates for HTS magnets can be an order of magnitude larger and take months to stabilize to within specified performance.(*15*) For instance, the magnetic field of the 1.1 GHz NMR spectrometer at NMRFAM drifted up rapidly after energization (50 Hz/hour) and reached a maximum approximately 6 months after installation (Fig. 2A). The drift then accelerated downward; after 10 months, the drift rate for ^13^C was measured at -9.6 ppb/hour (Fig. 2B, blue). This field drift could be compensated upon activating the ^2^H lock, reducing the variations over an 8-hour period by almost two orders of magnitude, from almost 80 ppb to less than 2 ppb (Fig. 2B, red). Although some fraction of this improvement may be achieved by a passive drift correction, this presumes a constant overall drift rate, which is not true for periods longer than few hours.

**Fig. 2.**
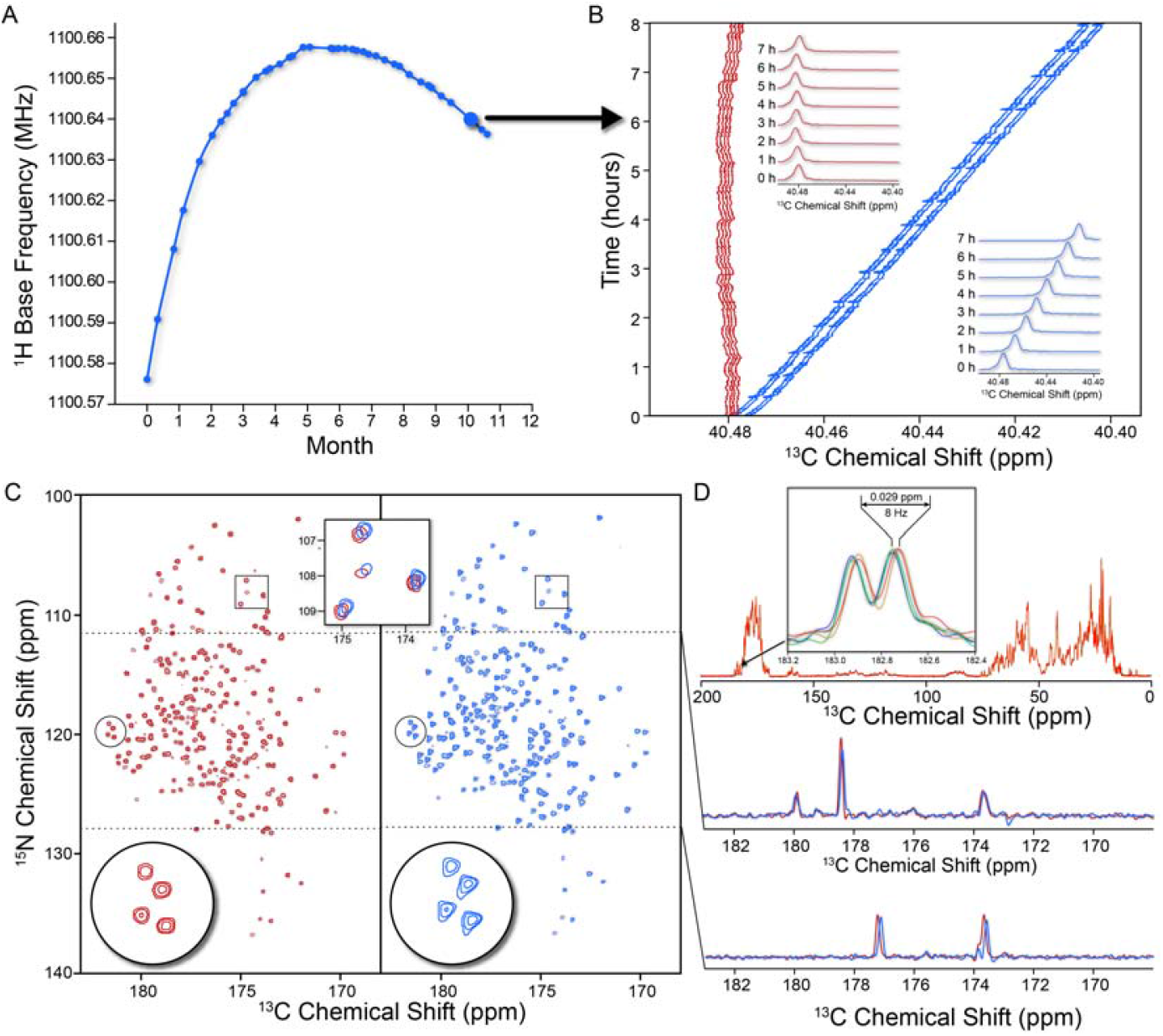
Magnetic field drift and the performance of external ^2^H lock. (A) Long-term post-installation field drift of the 1.1 GHz NMR spectrometer at NMRFAM. (B) Field drifts over 8 hours monitored using the downfield peak of adamantane with ^2^H lock on (red) and off (blue). The measurement was conducted ∼10 months after installation. (C) Comparison of ^15^N/^13^CO 2D correlation spectra of ToHo-1 β-lactamase acquired with ^2^H lock on (red), and off (blue). The LOW-BASHD was applied to both experiments. (D) Evaluation of long-term ^2^H lock performance using ^13^C 1D spectra of ToHo-1 β-lactamase acquired over 7 days.

To demonstrate the practical significance of the ^2^H lock for GHz-class spectrometers, we collected ^15^N/^13^CO 2D correlation spectra over 14 hours for Toho-1 β-lactamase, a 29 kDa bacterial enzyme involved in antimicrobial resistance,(*24*) with and without ^2^H lock using the 1.1 GHz NMR (Fig. 2C). We observed severe peak shape distortion without the ^2^H lock, with a notable shift in peak positions. These adverse effects are unacceptable for the acquisition of high-quality 2D or 3D datasets for macromolecules that may require several consecutive days of data collection. Uncertainties in peak positions also complicate procedures for resonance assignments, which are required for all subsequent stages of data analysis when studying structure and dynamics by NMR. In contrast, the ^2^H lock improved the sensitivity and eliminated artifactual peak asymmetries and referencing offsets (Fig. 2C and Table S1).(*25, 26*) The benefits of ^2^H lock are compounded for longer experiments; for example, with lock on, spectra varied by less than 0.03 ppm over 7 days (Fig. 2D), a ∼50-fold reduction relative to the native drift of the magnet (∼1.6 ppm).

### Temperature effect on external ^2^H lock

Another unique and challenging feature of HTS magnets that we observed is that field stability is highly sensitive to temperature variations in the magnet bore. These variations can arise from changes in experimental temperature, sample replacement, or probe reinsertion. We observed that re-establishing thermal equilibrium within the probe-magnet system after such changes could take several hours, consistent with previously reported stabilization times for GHz-class NMR systems.(*27*) Since all SSNMR probes thus far designed for these magnets require lowering the probe out of the magnet to change the sample, it is therefore essential to access the impact of temperature fluctuations on the performance of the external ^2^H lock. We first monitored the shift in ^13^C and ^2^H peak upon changing the experimental temperature through the variable temperature (VT) unit of the spectrometer. For the Black Fox probe, ^13^C and ^2^H signals moved in opposite directions due to field gradient variation (Fig. 3A). This causes the magnitude of change in ^13^C signal to be exacerbated due to temperature variations when the lock is engaged (Fig. 3B). We repeated the same measurement using CD_3_CN as the lock sample, which has an opposite and little response to temperature changes compared to D_2_O (Fig. S2). However, a similar drift trend was observed (Fig. 3C and 3D). This result indicates that the temperature-dependent peak position drift is predominantly driven by the inhomogeneous response of the field gradient of the HTS magnet to temperature changes. This effect is independent of the specific probe used, as a similar temperature effect was also observed for the Phoenix probe (Fig. 3E and 3F), though with a smaller amplitude and faster equilibrium due to the closer proximity of the coil to the magnetic field center. It is important to note that temperature-induced peak shifts are reversible once the thermal equilibrium is restored. Besides temperature, cryogen filling and powering the Bruker Smart Nitrogen Liquefier (BSNL) system on or off were also found to affect ^2^H lock performance significantly (Fig. S3).

**Fig. 3.**
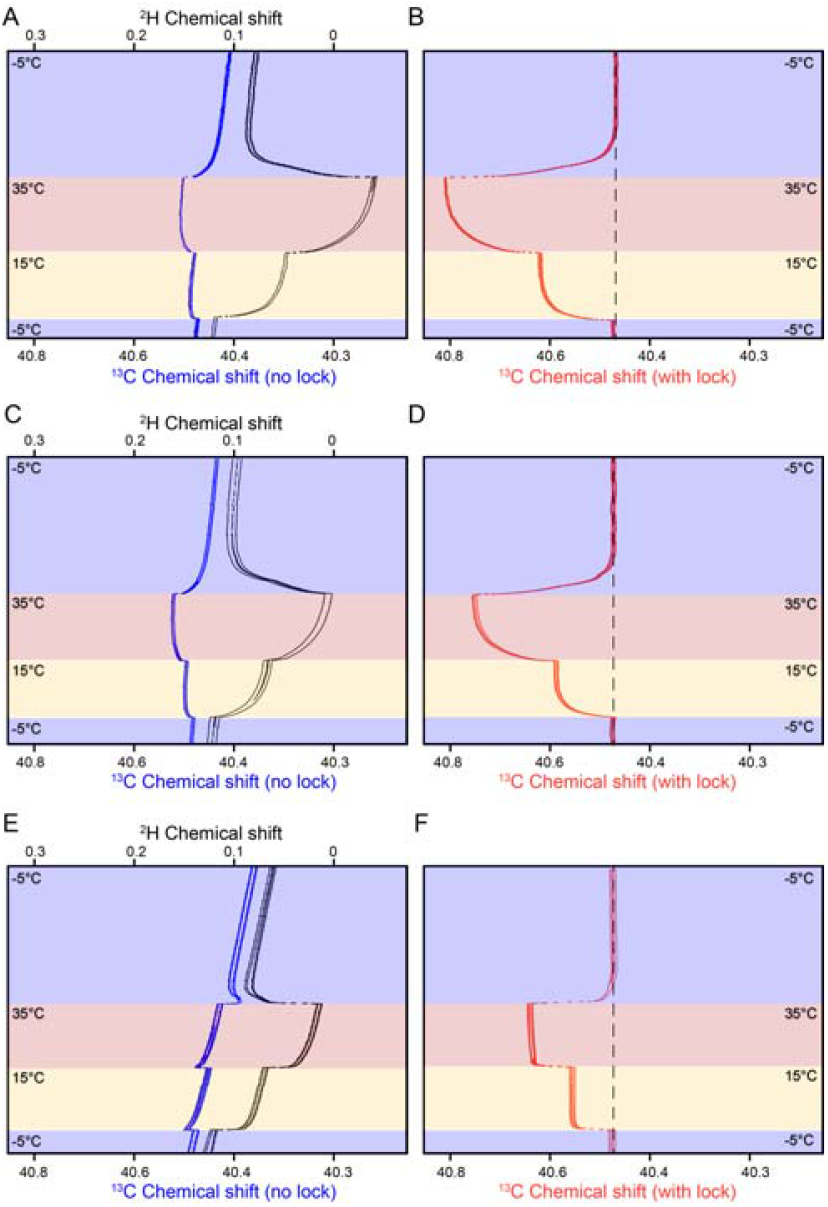
Temperature effect on ^2^H lock performance. (A) Temperature effects on downfield peak of adamantane in the sample coil (blue) and ^2^H peak of D_2_O in the lock coil (black) with ^2^H lock off. (B) Temperature effects on downfield peak of adamantane with ^2^H lock on. Measurements were conducted using the Black Fox 1.6 mm probe. The temperatures labeled in the figure represent the VT temperatures. (C) and (D), similar measurements as described in (A) and (B), but using CD_3_CN as lock solvent. (E) and (F), similar measurement as described in (A) and (B), but using Phoenix 1.6 mm probe. The VT temperature changes are virtualized by colors in the plots. The dashed lines in the right panels indicate that the chemical shift changes are reversible upon temperature changes.

### LOW-BASHD decoupling

The maximum spectral resolution achievable in SSNMR is determined almost entirely by the intrinsic properties of the sample itself, and this is true both at moderate magnetic field and at field above 1 GHz.(*28, 29*) For well-ordered microcrystalline protein samples, this allows, for example, the observation of fine structure due to the homonuclear *J*-couplings. With our instrument, the *J*-couplings are the dominant source of inhomogeneous line broadening and effective limit to the overall linewidth. Homonuclear decoupling is therefore essential to realizing the highest achievable resolution. Here we demonstrate that UHF is compatible with state-of-the-art homonuclear decoupling during direct detection, allowing remarkable resolution in protein SSNMR. Specifically, we make use of the LOW-BASHD approach to decouple the ∼55 Hz *J*_CαC’_ splitting during ^13^C detection.(*20*) Fig. 4A shows a comparison of ^15^N/^13^CO 2D correlation spectra of tryptophan synthase, an αββα heterodimer with an effective molecular weight of 72 kDa per αβ unit,(*30*) acquired with and without LOW-BASHD. The refocusing of *J*_CαC’_ during direct detection by LOW-BASHD results in an approximately 2-fold improvement in resolution and sensitivity (Fig.4A, Table S2). With 18 hours of data acquisition, approximately 500 unique well-resolved resonance peaks are observed in the 2D spectrum, consistent with the reported number of detectable residues.(*30*) Moreover, it substantially improves the alignment of resonance peaks in protein backbone assignments using 3D experiments (Fig. 4B).

**Fig. 4.**
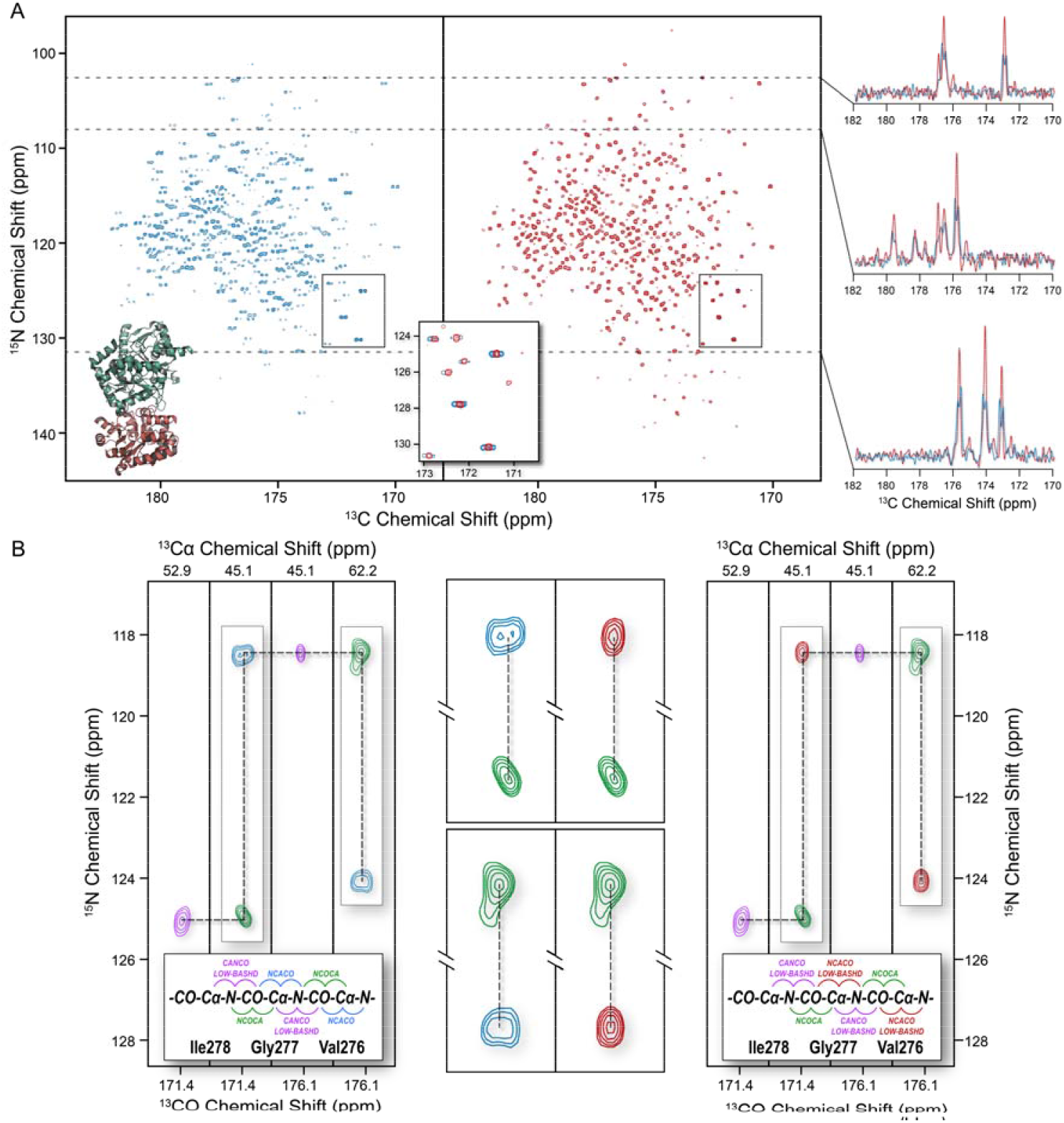
The LOW-BASHD approach improves sensitivity and resolution in ^13^CO detected experiments, facilitating the sequential assignments. (A) Comparison of ^15^N/^13^CO 2D correlation spectra of Tryptophan synthase acquired without LOW-BASHD (blue) and with LOW-BASHD (red). (B) Examples of 3D strip plots of tryptophan synthase for sequential assignments. The left plots contain NCACO (blue), NCOCA (green), and CANCO with LOW-BASHD (purple). For comparison, the right plots replace the NCACO to NCACO with LOW-BASHD (red). Middle panel compare the zoomed strips (highlighted in boxes) to illustrate the improved peak alignment achieved with LOW-BASHD.

## Discussion

The rapid field drift of HTS magnets has become a significant challenge, limiting the application of UHF NMR to complex research systems. Currently, the most widely used method for field drift correction in the SSNMR community is the passive drift compensation feature offered by Bruker Topspin software, which applies a constant linear correction to the magnetic field. While effective for slow and stable drift, this approach is less reliable for HTS magnets, where drift is fast and inherently nonlinear over extended periods, leading to cumulative errors. In contrast, the external ^2^H lock enables real-time correction of field fluctuations regardless of drift rate or linearity. The external ^2^H lock for SSNMR was first introduced by Paulson and Zilm in 2008,(*18*) using a spherical sample bulb filled with D_2_O solution placed above the probe and connected to a separate circuit from the top of the spectrometer. However, this design is incompatible with modern HTS-based GHz-class NMR spectrometers because of the short length of the sweet spot of the magnet, along with the design of the Bruker shim stack, which restricts the proximity of the lock coil to the sample coil. The probes presented in this study overcome these limitations by integrating the ^2^H lock coil directly into the probe head, minimizing the distance between the lock and sample coils to ensure compatibility with the restricted field range of HTS magnets.

The temperature-dependent shift of the ^2^H lock signal indicates that the external ^2^H lock cannot correct field disturbances caused by temperature or sample changes. Instead, the effect may be amplified due to temperature-dependent field gradient variations in the HTS magnet. In practice, 1D spectra should be collected regularly in the first few hours following a sample or temperature change to verify field stability. The time required to restore thermal equilibrium varies depending on the probe design. Probes with closer proximity between the sample and lock coils, and better thermal insulation design, tend to stabilize more quickly. For example, the Black Fox probe typically requires ∼6 hours to reach thermal equilibrium after temperature changes, whereas the Phoenix probe stabilizes within ∼2 hours. This difference is attributed to the proximity of the Phoenix lock coil to the magic-angle spinning stator, as well as a plastic coating layer inside the probe head cover that improves the thermal insulation. However, once thermal equilibrium is reached, the performance of the ^2^H lock is consistent across different probes. It is also important to note that temperature-induced peak shifts are fully reversible once the field stabilizes. Therefore, the ^2^H lock and spectral referencing must be conducted at a consistent temperature for upcoming research samples to obtain accurate chemical shift measurements.

The effectiveness of LOW-BASHD is determined by the magnitude of the *J*-coupling relative to other contributions to the linewidth through the effective ^13^C transverse relaxation time (*T’*_*2*_). Consequently, the resolution enhancement from LOW-BASHD will be less pronounced for signals where linewidths are predominantly determined by other homogeneous and inhomogeneous contributions. This explains in part why LOW-BASHD approach, originally developed for SSNMR, has been used primarily for ^13^C-detected experiments in protein solution NMR.(*31*) To maximize the ^13^C resolution enhancement offered by LOW-BASHD in MAS SSNMR, it is crucial to minimize other linebroading contributions, primarily ^1^H/^13^C heteronuclear dipolar couplings. This requires the use of high-power ^1^H decoupling, which can lead to significant RF-induced sample heating when applied for extended durations. To mitigate the sample heating, the probes described in this study incorporate a dual resonator low electric field (“Low-E”) design, which effectively reduces the heat generation from the ^1^H channel.(*32, 33*) This capability is particularly critical for ^13^C detection with GHz-class NMR, where probe RF efficiency for ^1^H is relatively low and signal lifetimes are prolonged. For example, on the 1.1 GHz NMR spectrometer, the ^15^N and ^13^C_α_ *T*_*2*_ relaxation times of the tryptophan synthase sample were measured at 23 ms and 9 ms, respectively. As a result, the total acquisition times for multi-dimensional experiments typically exceed 50 ms with high-power ^1^H decoupling. The Low-E resonator design is therefore essential for enabling extended high-power ^1^H decoupling while ensuring sample safety in ^13^C detected experiments using UHF NMR.

In summary, we introduced two SSNMR probes with distinct ^2^H lock coil designs, demonstrating the critical role of ^2^H lock in ensuring high-quality and accurate data acquisition on GHz-class NMR systems. The influence of temperature on field stability affects the ^2^H lock performance, underscoring the importance of maintaining consistent experimental conditions. Additionally, the exceptional resolution offered by UHF NMR exposes the effect of ^13^C-^13^C *J*-coupling splitting in ^13^C-detected experiments. This issue is effectively resolved through the application of the LOW-BASHD approach, which not only eliminates splitting but also substantially enhances sensitivity and ensures precise resonance assignments. To demonstrate the power of these techniques, we showcased an ultra-high resolution 2D spectrum of the 144 kDa protein tryptophan synthase, achieving unique site resolution for several hundred residues. We envision that the application of external ^2^H lock and LOW-BASHD in GHz-class NMR will open new opportunities in drug discovery, structural biology, and advanced materials research.

## Materials and Methods

### Instrumentation

All NMR experiments were performed on a Bruker NEO 1.1 GHz spectrometer at NMRFAM. Two probes assessed in this study were the Black Fox 1.6 mm HCN triple-resonance probe (Black Fox LLC) and the Phoenix 1.6 mm HXY triple-resonance probe (PhoenixNMR LLC). Both probes featured external ^2^H lock functionality, utilizing D_2_O sealed in a 1.5 mm OD × 12.5 mm long glass capillary (New Era) as lock sample. 20 mM Cu^2+^ doped into D_2_O to accelerate the ^2^H *T*_*1*_ relaxation. The capillaries were flame-sealed at one end. D_2_O was added via centrifuge until full, the other end was then sealed with epoxy (J-B WELD). Bubbles need to be avoided to achieve good shim and lock results. For the Black Fox probe, the capillary was inserted horizontally into the lock coil and fixed using two 1.2×0.6 mm O-rings. For the Phoenix probe, the capillary was inserted vertically into the lock sample slot, and the insertion hole was covered with a small piece of tape for stabilization. For ToHo-1 β lactamase and tryptophan synthase samples, all spectra were collected at 25 kHz MAS rate, and the VT temperature was set as -5 ºC. All spectra were referenced to DSS, using adamantane as a secondary external standard. The downfield ^13^C signal of adamantane was referenced to 40.48 ppm.

### Preparation of Toho-1 β-lactamase

Uniformly labeled ^13^C-^15^N Toho-1 was produced, purified, and crystallized following a modified version of a previously described protocol.(*24, 34*) In brief, BL21 *E. coli* cells transformed with a Toho-1 β-lactamase expression plasmid were overexpressed in 500 mL of kanamycin-supplemented minimal media containing 4.0 g ^13^C-glucose, 3.0 g ^15^N-NH_4_Cl, 200 mM Sorbitol, and 5 mM β-iene induced by 1 mM IPTG and grown overnight. Cells were harvested by centrifugation and resuspended in 20 mM MES, pH 6.5 (Buffer A) and lysed via sonication using a Branson 450D Digital Sonifier (Emerson Industrial Automation, St. Louis, MO, USA). Protein was purified through a 5 mL HiTrap SP Sepharose FF column (GE Healthcare, Pittsburgh, PA, USA) pre-equilibrated with Buffer A, and protein separation was achieved using a linear gradient with 20 mM MES, 300 mM NaCl, pH 6.5 (Buffer B) and protein elution monitored by UV absorbance. Collected protein fractions were pooled for size-exclusion chromatography using a 120 mL HiLoad Superdex S-200 column equilibrated with Buffer A. Elution was again tracked by UV absorbance, and the pooled protein fractions were concentrated using a 10 kDa MWCO Ultra Centrifugal Filter (Amicon) to a final concentration of 300 - 400 μM. Crystallization was initiated by mixing concentrated protein at a 1:1 ratio with crystal buffer (7 mM spermine, 30% PEG-8k) and incubated at 4 °C to allow crystallization over several days.

### Preparation of tryptophan synthase

Uniformly labeled ^13^C-^15^N Salmonelia typhimurium tryptophan synthase in *E. coli* was expressed and purified as previously described.(*35, 36*) Microcrystalline protein samples were prepared in 50 mM Cs-bicine, pH 7.8 crystal buffer containing 14% PEG-8000 and 3.0 mM spermine as previously described.(*35*)

### Mapping effective magnetic field range of the 1.1 GHz spectrometer

Field mapping was performed using the Phoenix 1.6 mm probe which was mounted on a probe frame attached to the Bruker shim stack, allowing free adjustment of the probe’s vertical position. A 1.6 mm rotor filled with D_2_O, spinning at 5.8 kHz, was used as the sample. All shim parameters were set to zero. The probe’s initial position was set to the highest allowable point, defined by the top of the shim stack. A ^2^H 1D spectrum was collected at this position, showing a coil image. The probe was then lowered in 5 mm increments, with ^2^H 1D spectra acquired at each step. The spectrum with the narrowest ^2^H peak suggested that the sample was near the magnetic field center. The lower limit of the effective field range was determined when the ^2^H signal became significantly broadened and undetectable.

### Shimming for double sample and ^2^H lock coils simultaneously

A dual-acquisition pulse sequence was employed to shim both coils simultaneously, combining a ^13^C one-pulse experiment with low-power ^1^H decoupling to acquire the ^13^C signal from adamantane and a ^2^H one-pulse experiment to detect the ^2^H signal from D_2_O. The adamantane sample, packed in a 1.6mm Varian style rotor, was spun at 30 kHz. OPTO was used to evaluate the effect of each shim parameter on both ^13^C and ^2^H signals to determine which parameters significantly influence each coil.(*22*) The information provided a guideline for manual adjustments. The ^13^C signal from the sample coil was shimmed first to achieve the narrowest possible ^13^C linewidth. Subsequently, shim parameters that had minimal impact on the sample coil but strongly affected the lock coil were adjusted to refine the ^2^H signal. The objective was to achieve a ^2^H linewidth below 20 Hz without noticeable splitting, while minimally compromising the ^13^C linewidth. Once optimal shimming was achieved, the shim parameters were saved. For future shimming, only significant adjustments to the z value are typically required, while other parameters usually need minor fine-tuning only.

### Measuring ^13^C and ^2^H resonance peak drifting trajectories

A ^13^C one-pulse experiment with low-power ^1^H decoupling was used to record the adamantane ^13^C resonance peak drift. A 2.5µs 90º pulse was applied on ^13^C channel, and signals were acquired with 15 kH continuous wave (CW) ^1^H decoupling. The total acquisition time was 500 ms. To track ^2^H signal drift, the lock function was turned off. A ^2^H one-pulse experiment, using a 20 µs 90º pulse, was used to generate the ^2^H signal from the lock channel, with an acquisition time of 500 ms. The ^13^C and ^2^H signals were acquired simultaneously using the same dual acquisition pulse program. The 1D spectra were recorded at 30 s intervals, and the data were stored in pseudo-2D containers for further processing. For the ^13^C peak drifting measurement with ^2^H lock, the same ^13^C 1D spectra were acquired as described above except using a regular single ^13^C acquisition pulse program.

### ^2^H lock setup for research samples

After completing the shimming process, the stability of the magnetic field was assessed by tracking the ^13^C signal of adamantane with the ^2^H lock activated. Once the ^13^C peak position showed no significant drift, the field value was adjusted to position the left peak of adamantane at 40.48 ppm, ensuring an accurate reference to DSS. Sample changes do not require deactivating the ^2^H lock, as the lock function automatically reactivates once the probe is back to its previous position in the magnet. Note that taking probe in and out may disturb the magnetic field, so additional time is needed for the field to stabilize following the sample change, typically 2 to 6 hours. During this period, pulse sequences can be optimized. For consistent and accurate chemical shift referencing for research samples, both the adamantane calibration and the experiments on research samples must be conducted at the same temperature.

### ^15^N/^13^CO 2D correlation spectra for ToHo-1 β lactamase and Tryptophan synthase

The 2D NCO experiment was performed with the ^13^C carrier frequency set at 175 ppm. The ^15^N polarization was prepared by an adiabatic CP with a downward tangential ramp pulse on the ^1^H channel, with a contact time of 2.0 ms, ^15^N RF amplitude of 40 kHz, and an average ^1^H RF amplitude of 60 kHz. ^15^N polarization was then transferred to ^13^CO using CP with an upward tangential ramp on the ^13^C channel, with a contact time of 7.0 ms, ^15^N RF amplitude of 10 kHz, and an average ^13^C RF amplitude of 16 kHz. A CW decoupling of ^1^H at 100 kHz was applied during this CP period. The t_1_ acquisition time was 41.0 ms with a 40 µs increment for 1024 complex points. A 5.2 µs ^13^C π-pulse was applied at the center of the t_1_ period to decouple the *J*_*15N-13C*_. The t_2_ acquisition time was 30.7 ms with a 5 µs dwell time for 3072 complex points. SPINAL-64 ^1^H decoupling at 100 kHz was applied during acquisition. For LOW-BASHD experiments, LOW-BASHD detection was implemented in the direct dimension to decouple the ^*1*^*J*_*C*α*-C’*_ using τ_Dec_ = 3.2 ms and 72.5 µs cosine modulated Gaussian π-pulses, modulated at 37 kHz.(*20*) The recycle delay was 1.5 s. The total experimental time was 14 hours.

### ^15^N/^13^C_α_/^13^CO 3D correlation spectrum (NCACO) for Tryptophan synthase

The 3D NCACO experiment was performed with the ^13^C carrier frequency set at 55 ppm. The ^15^N polarization was prepared by an adiabatic CP with a downward tangential ramp pulse on the ^1^H channel, with a contact time of 1.5 ms, ^15^N RF amplitude of 40 kHz, and an average ^1^H RF amplitude of 59 kHz. ^15^N polarization was then transferred to ^13^C_α_ using CP with an upward tangential ramp on the ^13^C channel, with a contact time of 7.0 ms, ^15^N RF amplitude of 10 kHz, and an average ^13^C RF amplitude of 15 kHz. A CW decoupling of ^1^H at 100 kHz was applied during this CP period. The ^13^C_α_/^13^CO polarization transfer was achieved using a 100.0 ms CORD mixing period with a ^1^H RF amplitude of 25 kHz.(*37*) The t_1_ acquisition time was 15.4 ms with a 120 µs increment for 128 complex points. A 5.2 µs ^13^C π-pulse was applied at the center of the t_1_ period to decouple the *J*_*15N-13C*_. The t_2_ acquisition time was 6.4 ms with an 80 µs increment for 80 complex points. A 5.2 µs ^13^C hard π-pulse, a 300 µs ^13^C soft π-pulse with RSNOB shape at 55 ppm, and a 14.8 µs ^15^N π-pulse were applied at the center of the t_2_ period to decouple the *J*_*13C*_ *-*_*13CX*_ and *J*_*13C*α*-15N*_. The t_3_ acquisition time was 30.7 ms with a 5 µs dwell time for 3072 complex points. SPINAL-64 ^1^H decoupling at 100 kHz was applied during acquisition. For LOW-BASHD version of NCACO experiment, LOW-BASHD was implemented in the direct dimension to decouple the ^1^*J*_*C*_α*-C’*__ using τ_dec_ =3.2 ms and 72.5 µs cosine modulated Gaussian π-pulses, modulated at 37 kHz. The recycle delay was 1.5 s. The 3D spectrum was acquired using a 25% NUS schedule and the total experimental time was 113.4 hours.

### ^15^N/^13^CO/^13^C_α_ 3D correlation spectrum (NCOCA) for Tryptophan synthase

The 3D NCOCA experiment was performed with the ^13^C carrier frequency set at 175 ppm. The ^1^H to ^15^N CP and the ^15^N to ^13^CO CP transfer conditions were identical to the 2D NCO experiment as described above. The ^13^CO_α_/^13^C_α_ polarization transfer was achieved using a 100.0 ms CORD mixing period with a ^1^H RF amplitude of 25 kHz. The t_1_ acquisition time was 15.4 ms with a 120 µs increment for 128 complex points. A 5.2 µs ^13^C π-pulse was applied at the center of the t_1_ period to decouple the *J*_*15N-13C*_. The t_2_ acquisition time was 7.7 ms with a 160 µs increment for 48 complex points. A 5.2 µs ^13^C hard π-pulse, a 300 µs ^13^C soft π-pulse with Rsnob shape at 175 ppm, and a 14.8 µs ^15^N π-pulse were applied at the center of the t_2_ period to decouple the *J*_*13CO-13CX*_ and *J*_*13CO-15N*_. The t_3_ acquisition time was 30.7 ms with a 5 µs dwell time for 3072 complex points. SPINAL-64 ^1^H decoupling at 100 kHz was applied during acquisition. The recycle delay was 1.5 s. The 3D spectrum was acquired using a 25% NUS schedule and the total experimental time was 136.2 hours.

### ^13^C_α_/^15^N/^13^CO 3D correlation spectrum (CANCO) for Tryptophan synthase

The 3D CANCO experiment was performed with the ^13^C carrier frequency set at 175 ppm. ^13^C_α_ polarization was prepared via adiabatic CP using a downward tangential ramp pulse on the ^1^H channel, with the ^13^C frequency adjusted to 55 ppm for ^1^H-^13^C_α_ transfer. The CP contact time was 1.5 ms, ^13^C RF amplitude was 108 kHz, and the average ^1^H RF amplitude was 82 kHz. Polarization was then transferred from ^13^C_α_ to ^15^N using a CP with an upward tangential ramp on the ^13^C channel. The contact time was 7.0 ms, ^15^N RF amplitude was 10 kHz, and the average ^13^C RF amplitude was 16 kHz. A 100 kHz ^1^H CW decoupling was applied during this CP period. The ^15^N polarization was transferred to ^13^CO using CP with an upward tangential ramp on the ^13^C channel. The ^13^C frequency was changed back to 175 ppm before this CP. The contact time was 7.0 ms, with a ^15^N RF amplitude of 10 kHz, and an average ^13^C RF amplitude of 17 kHz. A 100 kHz ^1^H CW decoupling was applied during this CP period. The t_1_ acquisition time was 6.4 ms with an 80 µs increment for 80 complex points. A 5.2 µs ^13^C hard π-pulse and a 300 µs ^13^C soft π-pulse with Rsnob shape at 55 ppm were applied at the center of the t_1_ period to decouple the *J*_*C*α*- CX*_. The t_2_ acquisition time was 15.4 ms with a 120 µs increment for 128 complex points. A 5.2 µs ^13^C π-pulse was applied at the center of the t_2_ period to decouple the *J*_*N-C*_. The t_3_ acquisition time was 30.7 ms with a 10 µs dwell time for 1536 complex points. A SPINAL-64 ^1^H decoupling at 100 kHz was used during acquisition. Additionally, LOW-BASHD detection was implemented in the direct dimension to decouple the *J*_*C*α*-CO*_ using τ_dec_=3.2 ms and 72.5 µs Gaussian π-pulses, modulated at 37 kHz. The recycle delay was 1.5 s. The 3D spectrum was acquired using a 25% NUS schedule and the total experimental time was 107.4 hours.

## Author contributions

Conceptualization: SW, CMR. Methodology: SW, TR, JAS, PLG, CMR. Investigation: SW, TR, CGW, OAW, AFT, LJM, CMR. Visualization: SW. Supervision: CMR, LJM. Writing—original draft: SW, CMR, LJM. Writing—review & editing: SW, TR, JAS, PLG, OAW, CGW, AFT, LJM, CMR

## Data and materials availability

All data needed to evaluate the conclusions in the paper are present in the paper and/or the Supplementary Information. The raw NMR data is available on the website of Network of Advanced NMR. The pulse sequences with LOW-BASHD decoupling were developed for the Bruker NEO console using Topspin 4.4.0. The pulse programs are available upon request.

## Supplementary Materials

