## Supporting Information for "Ultra-High Resolution Solid-State NMR for High Molecular Weight Proteins on GHz-Class Spectrometers"

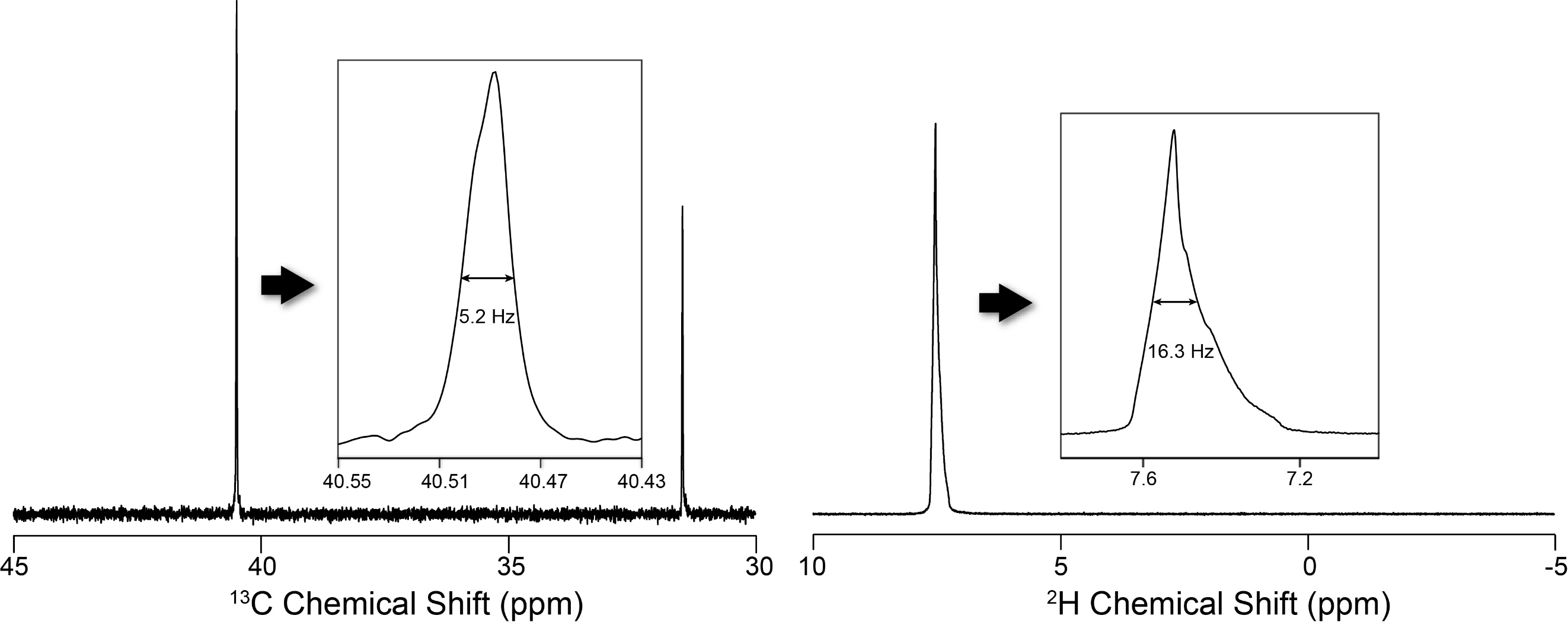

**Fig. S1. The best shimming result for Phoenix 1.6mm probe.** The left panel represents the ^13^C signal of the adamantane in the sample coil, while the right panel represents the ^2^H signal of D_2_O from the lock coil. The ^13^C and ^2^H signals were shimmed simultaneously using a dual acquisition pulse program.

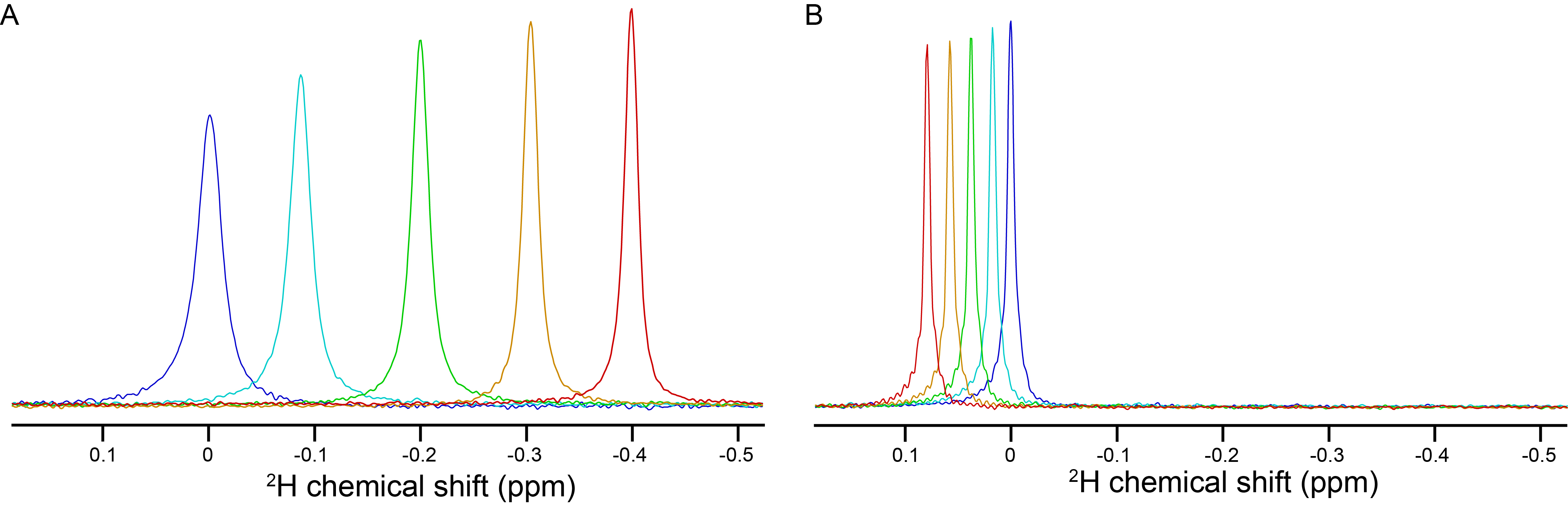

**Fig. S2. Comparison of temperature-dependent ^2^H chemical shift changes of D_2_O and CD_3_CN.** (A), The ^2^H 1D spectra of D_2_O acquired at 2 ºC (blue), 12 ºC (cyan), 22 ºC (green), 32 ºC (yellow), and 42 ºC (red). (B), The ^2^H 1D spectra of CD_3_CN acquired at 2 ºC (blue), 12 ºC (cyan), 22 ºC (green), 32 ºC (yellow), and 42 ºC (red). D_2_O and CD_3_CN have opposite response upon temperature change, and the amplitude for CD_3_CN is 5-fold less than D_2_O. The datasets were acquired using a 500 MHz solution NMR.

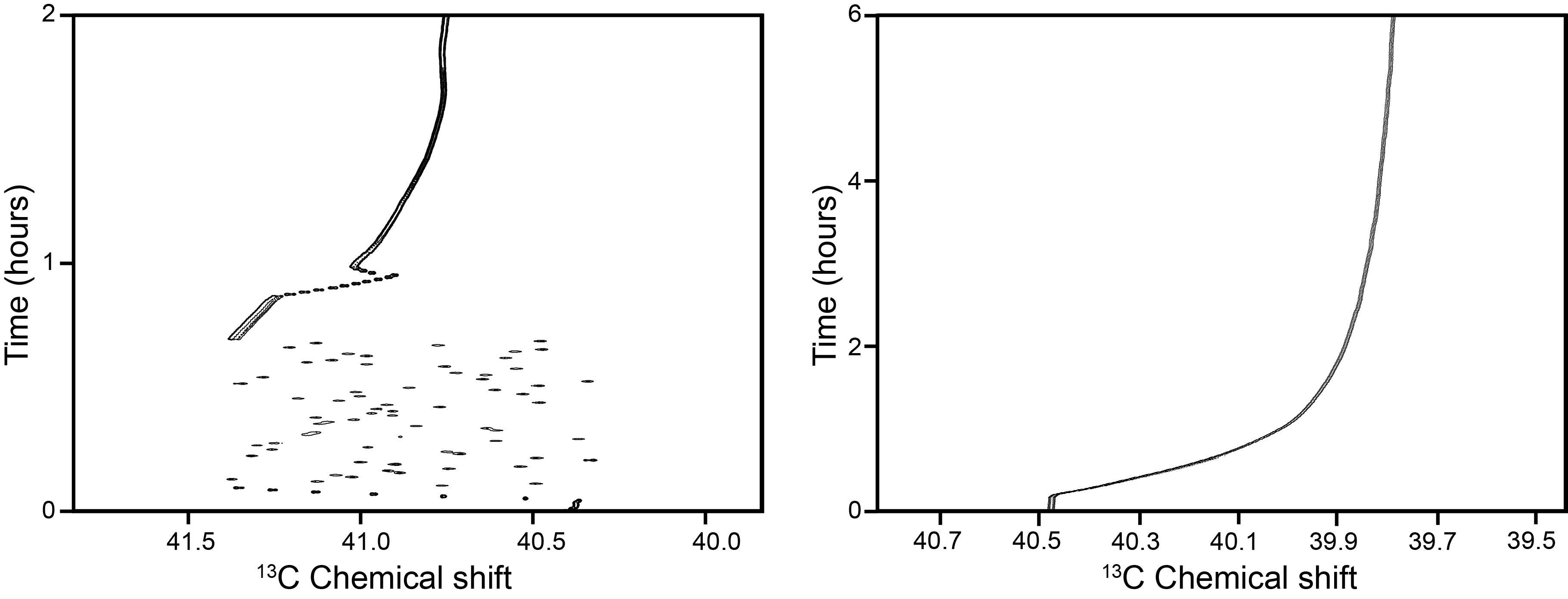

**Fig. S3. Impact of liquid Helium filling and power BSNL on/off on the ^2^H lock performance.** The shift of the downfield ^13^C peak of adamantane was recorded in these measurements. The liquid helium filling significantly disturbs the magnetic field stability, which makes the peak move discreetly in a 1 ppm range (left panel). Switching on BSNL causes a systematic upfield drift of the peak at approximately 0.7 ppm. And this effect took approximately 6 hours to be stabilized (right panel).

**Table S1.** Sensitivity and resolution comparison for ^15^N/^13^CO correlation 2D spectra of ToHo-1 β-lactamase with and without the ^2^H lock ^a^

|  | ^2^H lock on | | ^2^H lock off | |
| --- | --- | --- | --- | --- |
|  | Average ^b^ | SD ^c^ | Average | SD |
| Signal-to Noise Ratio (SNR) | 46.0 | 21.3 | 42.0 | 20.2 |
| ^15^N linewidth (Hz) | 25.7 | 4.3 | 27.0 | 4.5 |
| ^13^CO linewidth (Hz) | 41.0 | 6.1 | 43.5 | 7.9 |

a. LOW-BASHD was applied to both experiments, and identical processing parameters were used.

b. Average spectral SNR and linewidth values were calculated from the 240 strongest peaks in each spectrum.

c. SD represents the variation in SNR or linewidth among the 240 selected peaks, indicating the spread of values around the average.

**Table S2.** Sensitivity and resolution comparison for ^15^N/^13^CO correlation 2D spectra of Tryptophan synthase with and without LOW-BASHD ^a^

|  | With LOW-BASHD | | Without LOW-BASHD | |
| --- | --- | --- | --- | --- |
|  | Average | SD ^b^ | Average | SD |
| Signal-to Noise Ratio (SNR) | 24.0 | 9.4 | 12.7 | 5.8 |
| ^13^CO linewidth (Hz) ^c^ | 39.7 | 4.8 | 54.3 | 21.1 |

a. A spectral region (^15^N: 122.1–127.0 ppm, ^13^C: 179.7–182.2 ppm) containing only isolated peaks was selected for SNR and linewidth evaluation to minimize bias from peak overlap. The LOW-BASHD spectrum contains 15 resonance peaks in this region, while the spectrum without LOW-BASHD shows 30 peaks due to splitting. Both spectra were processed using identical parameters.

b. SD represents the variation in SNR or linewidth among the selected peaks, indicating the spread of values around the average.

c. The ^13^CO linewidth values listed do not account for peak splitting.
